# Abelson tyrosine-protein kinase 2 Regulates Myoblast Proliferation and Controls Muscle Fiber Length

**DOI:** 10.1101/159822

**Authors:** Jennifer Kim Lee, Steven J. Burden

## Abstract

Muscle fiber length is nearly uniform within a muscle but widely different among muscles. Here, we show that Abelson tyrosine-protein kinase 2 (Abl2) has a key role in regulating myofiber length, as a loss of Abl2 leads to excessively long myofibers in the diaphragm and other muscles. Increased myofiber length is caused by enhanced myoblast proliferation, expanding the pool of available myoblasts and leading to increased myoblast fusion. Abl2 acts in myoblasts, but expansion of the diaphragm muscle causes a reduction in size of the diaphragm central tendon. Replacement of tendon for muscle is likely responsible for the reduced stamina of *abl2* mutant mice. Further, ectopic muscle islands, each composed of myofibers of uniform length and orientation, form within the central tendon in *abl2*^*+/-*^ mice. Induction of *scleraxis* in tendon cells at the ends of these muscle islands suggests that myofibers stimulate tendon differentiation, which in turn regulates myofiber length.

## Introduction

Skeletal muscle fibers display a wide diversity in size both within individual muscles and among different muscles. This heterogeneity in muscle fiber diameter is initiated during development and regulated throughout life, as muscle fibers adapt their size in response to metabolic demands and neural activity, prompting muscle growth or leading to muscle atrophy (Schiaffino et al 2013, Glass 2003). Although the mechanisms that control muscle fiber diameter are reasonably well-understood (Schiaffino et al 2013, Glass 2003), the mechanisms that regulate muscle fiber length and ensure a nearly common myofiber length within a muscle, but a wide diversity in length among different muscles, are largely unknown. In principle, the mechanisms for setting muscle fiber length might be regulated by the size of the myoblast pool that is available to fuse, the propensity of myoblasts to fuse, the available space that is set by the positions of fixed skeletal elements, or all of these mechanisms.

Even less is known about how signaling between muscle and tendon cells influences muscle fiber growth, differentiation and orientation. Studies in *Drosophila* suggest that tendon-like cells, pre-positioned at the margins of a muscle, provide guidance cues that direct and orient myotube elongation as well as attachment sites for muscle (Frommer et al 1996, Schnorrer and Dickson 2004, Wayburn and Volk 2009). Although there is good evidence that muscle and tendon cells exchange signals to mutually control their differentiation in *Drosophila* (Wayburn and Volk 2009, Becker et al 1997, Volk and VijayRaghavan 1994, Schnorrer and Dickson 2004), far less is known about this process in vertebrates (Kardon 1998, Schweitzer et al 2010). As such, it remains possible that muscle fibers provide signals to one another (Ho et al 1983), influence the arrangement and properties of muscle interstitial cells (Mathew et al 2001) or alter the structure of the extracellular matrix between muscle fibers to ensure a common muscle fiber length and orientation (Hauschka and Konigsberg 1966).

Here, we show that a loss of Abelson related kinase 2 (Abl2), a non-receptor tyrosine kinase, selectively in myoblasts, leads to enhanced myoblast proliferation and an increase in muscle fiber length, consistent with the idea that the size of the myoblast pool has an important influence on muscle fiber length. As a consequence of muscle expansion, the size of tendon is reduced. Moreover, we show that ectopic muscle islands, surrounded by tendon cells, form in *abl2* heterozygous mice, yet the muscle fibers within these islands are of uniform length and orientation. These findings indicate that pre-positioned tendon cells are not essential to define the length and orientation of myofibers. Because specialized tendon cells form at the ends of these muscle islands, our results raise the possibility that a pioneering myotube induces tendon cells to organize and direct the orientation of later forming myotubes.

## Results

### Abl2 regulates muscle fiber length

The Abelson family of non-receptor tyrosine kinases, which includes Abl1 (c-abl) and Abl2 (also known as Arg), are widely expressed and crucial mediators of growth factor and adhesion receptors that regulate cell proliferation and cytoskeletal remodeling. Although *abl2*^-/-^ mice survive postnatally, *abl1*^-/-^ mice die at birth (Tybulewicz et al 1991). Thus, we studied *abl1*^-/-^ and *abl2*^-/-^ mice at E18.5, one day prior to birth. We began by examining the diaphragm muscle as the muscle can be readily viewed in its entirety as a whole-mount preparation, simplifying histological analysis and providing a comprehensive view of the muscle. Muscle fiber development appeared normal in *abl1*^-/-^ mice, as muscle fibers extend radially from the rib cage, converge medially around the central tendon, and attach to the central tendon (Hallock 2011). In the absence of Abl2, however, the diaphragm muscle fibers are excessively long, and the muscle nearly consumes the area normally occupied by the central tendon (Figure 1A). We measured muscle fiber length from insertion points at the ventral rib to the central tendon and found that muscle fibers in the diaphragm muscle are 1.7-fold longer in *abl2*^-/-^ mice than wild type littermate controls (Figure 1B). This expansion of muscle, evident during development, persists in adult *abl2* mutant mice. At all times, muscle length is enhanced while myofiber cross-sectional area remains normal (Figure S1).

**Figure 1.**
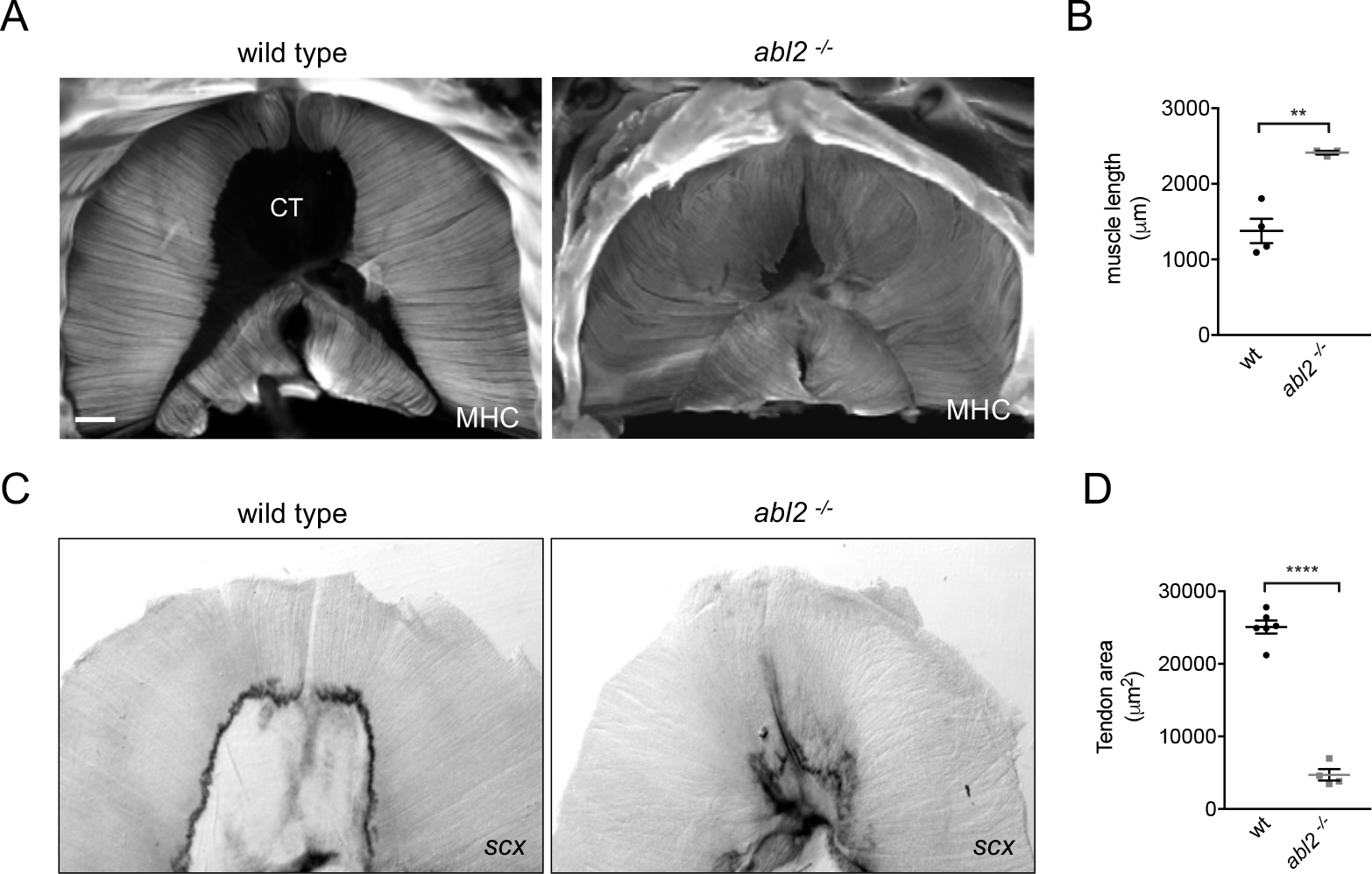
Muscle fibers are excessively long and the central tendon is reduced in size in *abl2* mutant mice. Whole mounts of muscle were stained with antibodies to myosin heavy chain (MHC). (A) Muscle fibers in the diaphragm muscle normally extend from the ribcage and attach medially to the central tendon (CT). Diaphragm muscle fibers in *abl2* mutant mice are extraordinarily long and the CT is diminished in size. (B) The mean length of the muscle, measured in the ventral quadrant of the diaphragm, is ∼1.7-fold longer in abl2 mutant than wild type mice. (C,D) The central tendon, circumscribed by high *scx* expression in tendon cells at the myotendinous junction, is reduced in size in *abl2* mutant mice. Scale bar is 500 μm in A.

### The size of the diaphragm central tendon is reduced in *abl2* mutant mice

To determine whether the elongated diaphragm muscle fibers grew over the normal central tendon domain or whether the central tendon was proportionally reduced in size we visualized the central tendon by using *in situ* hybridization to detect *scleraxis* (*scx*) expression. We found that *scx*, a transcription factor that is expressed in tendon precursor cells (Schweitzer et al 2001) is highly expressed in tendon cells at the myotendinous junction (MTJ) (Figure 1C). The area of the central tendon, circumscribed by *scx* expression, was reduced in *abl2*^-/-^ mice (Figure 1C). Thus, muscle fiber overgrowth and a reduction in size of the central tendon appear to be coordinated.

### Abl2 regulates muscle fiber length in several muscle groups

To determine whether Abl2 had a similar role in other muscles, we analyzed intercostal, levator auris longus and hind limb muscles in adult mice. We found that muscle fibers in intercostal muscles, which ordinarily extend from one rib to the adjacent rib, were likewise excessively long and extended over multiple ribs (Figure 2A). Similarly, muscle fibers in the levator auris longus muscle, which normally terminate at a midline tendon boundary, appeared wavy (Figure 2B), suggesting that some of the increased length was accommodated by bending of the myofiber (Figure 2B). To determine whether the midline tendon area was reduced in the levator auris longus muscle, we analyzed transgenic mice that express GFP under the control of the *scx* promoter in an *abl2*^-/-^ background (Schweitzer et al 2001, Pryce at al 2007). We found that the midline tendon domain, marked by GFP expression, was smaller in *abl2*^-/-^ than control mice. (Figure 2B). Thus, muscle fiber overgrowth and a reduction in size of the midline tendon appear to be coordinated in the levator auris longus muscle, similar to the diaphragm muscle. In contrast, we found no evidence for an increase in muscle fiber diameter length or cross-sectional area in limb muscles (Figure S2). Together, these data show that Abl2 has a role in regulating muscle fiber length in many but not all muscles.

**Figure 2.**
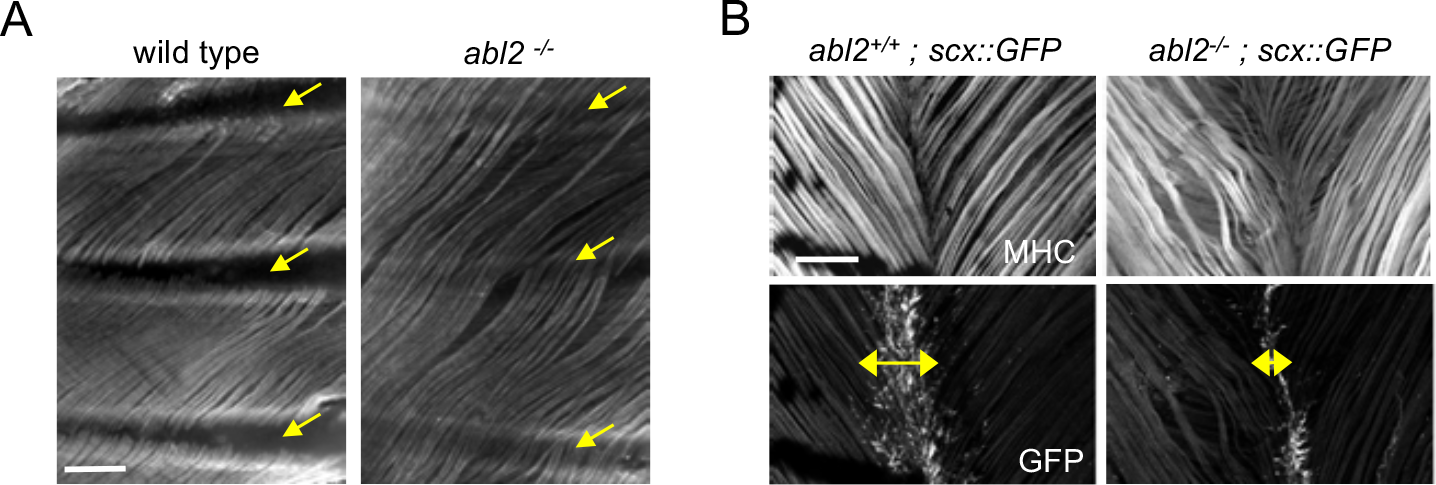
Muscle fibers are excessively long in intercostal and levator auris muscles in *abl2* mutant mice. Whole mounts of muscle were stained with antibodies to myosin heavy chain (MHC). (A) Intercostal muscle fibers normally extend from one rib to the adjacent rib (yellow arrows) but extend and cross over one or more ribs in *abl2* mutant mice. (B) Muscle fibers in the levator auris muscle appear wavy in *abl2* mutant mice. Moreover, the midline tendon, marked by *scx::GFP*, is reduced in size (double headed arrows). Scale bar is 250 μm in A and B.

### A loss of Abl2 leads to an increase in myoblast fusion

To determine whether the elongated muscle fibers arise from an increase in myoblast fusion or an increase in cytoplasmic volume without cell fusion, we counted the number of myofiber nuclei in diaphragm muscle fibers from E18.5 wild type and *abl2*^-/-^ mice. We counted myofiber nuclei in two ways: first, we dissociated muscle fibers and counted myonuclei in individual myofibers (Figure 3A,B); second, we measured myonuclei number in serial cross-sections of the diaphragm muscle, extending from the ribcage to the central tendon (Figure 3C,D). By both methods, we found that the increase in muscle fiber length was accompanied by a proportional increase in myonuclei number.

**Figure 3.**
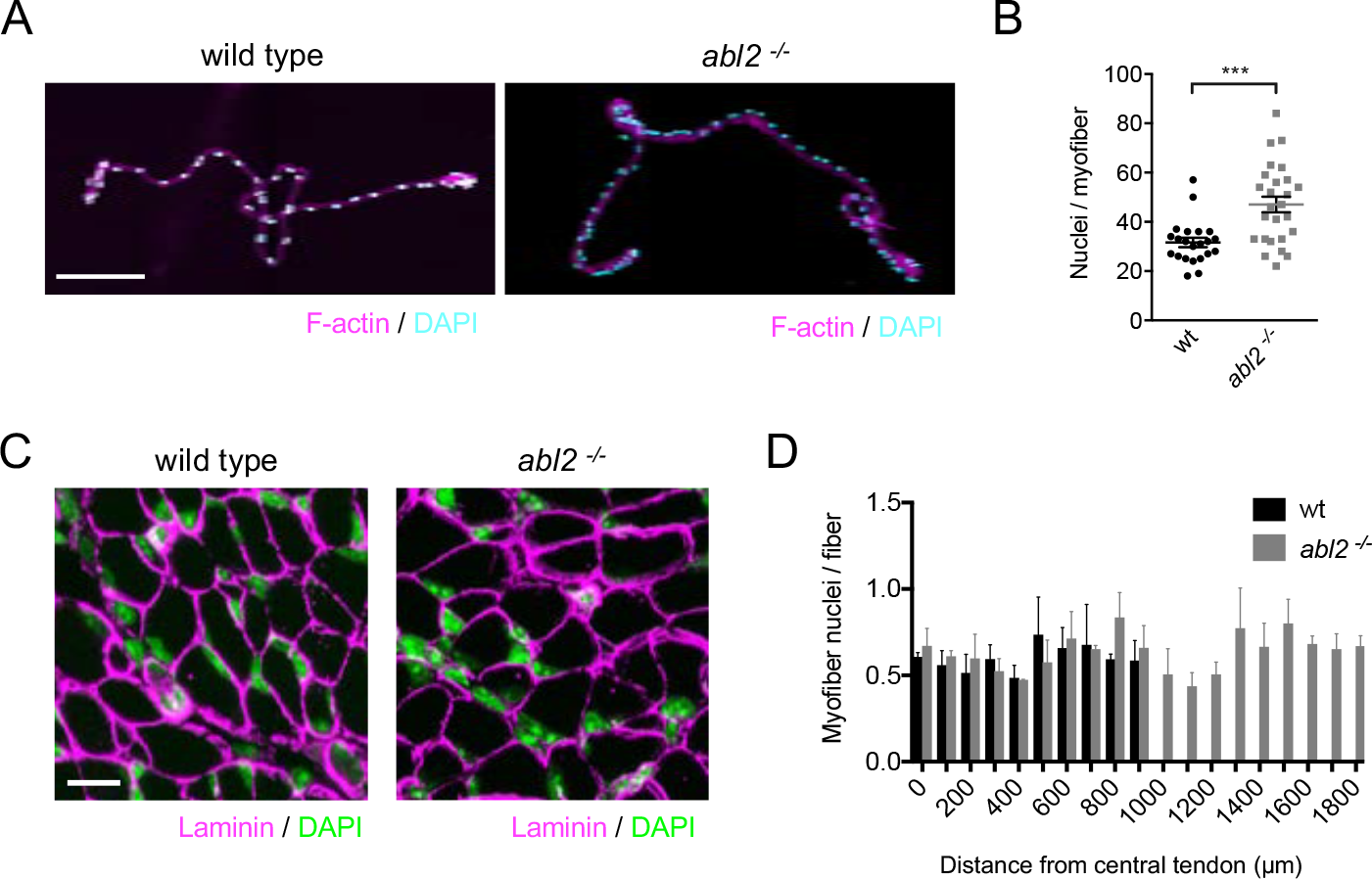
The increase in muscle fiber length is accompanied by an increase in myonuclei number. (A) Single fibers dissociated from the diaphragm muscle were stained with Phalloidin (F-actin) and DAPI. (B) The scatter plot shows the number of nuclei per myofiber for individual dissociated myofibers, dissected from 3 individual *abl2* mutant and 3 wild type mice, as well as the mean ± SEM. (C) Representative images from serial cross sections of the diaphragm muscle, stained for nuclei (DAPI) and with antibodies to Laminin, are shown here. (D) Myonuclei are similarly spaced along the entire length of muscle from wild type and *abl2* mutant mice. The graph shows the mean ± SEM per field of view in serial cross sections taken every 100 μm from the central tendon to the ribcage of 3 wild-type and 3 *abl2* mutant mice. Scale bar is 150 μm in A and 10 μm in C.

### Abl2 acts in myoblasts to control muscle fiber growth

We performed whole mount *in situ* hybridization experiments to determine where *abl2* is expressed in the diaphragm during development. We found that *abl2* is highly expressed in the diaphragm muscle and absent from the central tendon in E18.5 mice (Figure 4A,B), suggesting that *abl2* acts in myoblasts and/or myofibers to regulate muscle growth. To investigate whether Abl2 is expressed in myoblasts or multinucleated myotubes, we analyzed Abl2 protein expression during muscle differentiation in C2C12 cells. We found that Abl2 expression is high in myoblasts and declines during differentiation (Figure S3). These results indicate that myoblasts rather than myotubes are the major source of Abl2 expression in muscle.

**Figure 4.**
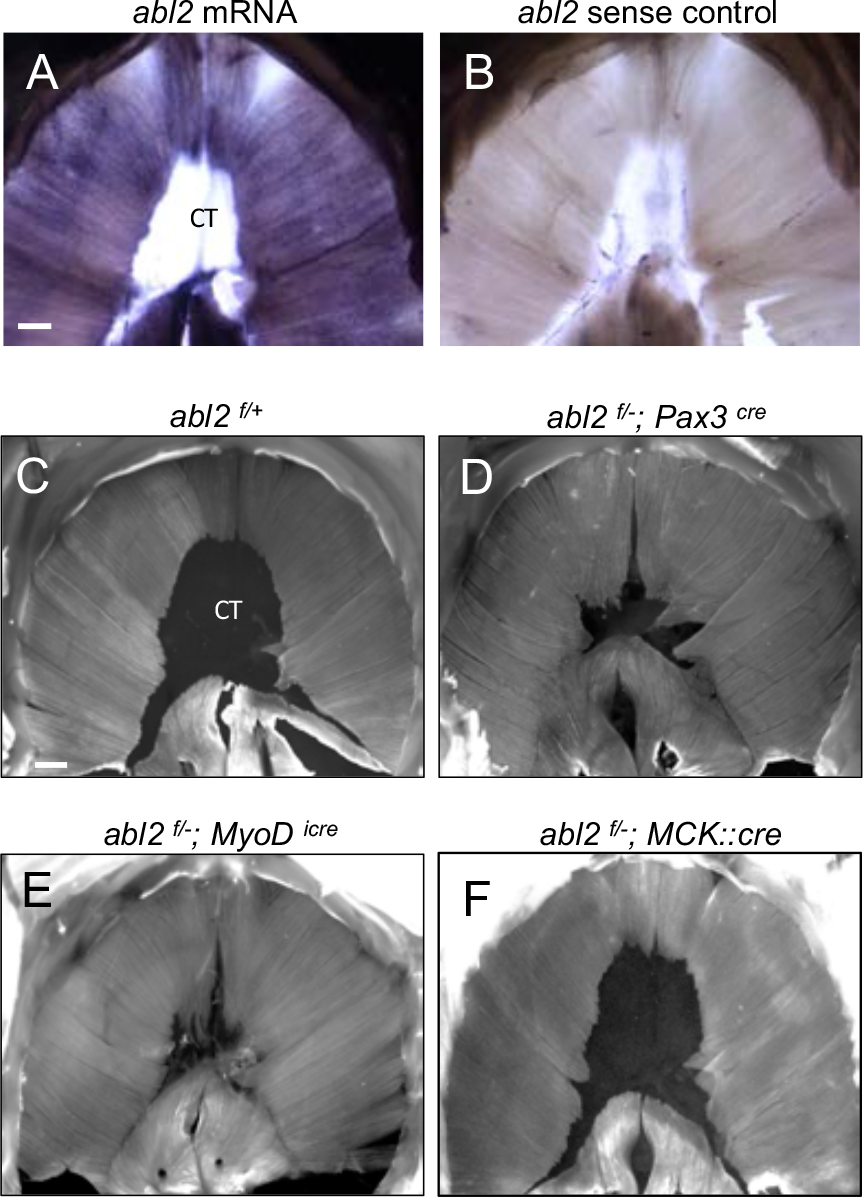
Abl2 acts in myoblasts to regulate muscle development. (A,B) *abl2 mRNA* is highly expressed in the muscle but not within the CT of the diaphragm. (C) Muscle and tendon development appear normal in *abl2*^*f*/+^ control mice. Muscle length is increased and tendon size is reduced by conditionally inactivating *abl2* in (D) muscle precursors (*abl2*^*f*/-^ *Pax3*^cre^), or (E) committed myoblasts (*abl2*^*f*/-;^ *MyoD*^*icre*^). (F) Muscle and tendon development appear normal by inactivating *abl2* in mature myotubes (*abl2*^*f*/-^; *MCK::cre*). Scale bars are 500 μm in A and C.

To examine the site of action of *abl2* during myogenesis, we obtained mice bearing an *abl2* allele with *loxP* sites and generated conditional *abl2* mutants using mice that express Cre recombinase in pre-migratory myogenic precursor cells (*Pax3^Cre^*), committed myoblasts (*MyoD^iCre^*) or multinucleated myofibers (*MCK::Cre*) (Engleka et al 2005, Relaix et al 2006, Kanisicak et al 2009, Braun et al 1994, Bruning et al 1998).

Selective inactivation of *abl2* in pre-migratory myogenic precursor cells (*abl2*^*f*/-^; *Pax3*^cre^) led to muscle defects like those found in *abl2*^-/-^ mice (Figure 4C,D). Likewise, inactivation of *abl2* in committed myoblasts (*abl2*^*f*/-^ *MyoD*^*icre*^) caused aberrations in muscle development identical to those found in *abl2*^-/-^ mice (Figure 4C,E). In contrast, inactivation of *abl2* in myofibers (*abl2*^*f*/-^ *MCK::Cre*) did not cause a muscle phenotype (Figure 4F). These findings indicate that Abl2 is required in myoblasts rather than multinucleated myotubes to regulate muscle and tendon growth.

### *Abl2* mutant mice display impaired motor endurance

Diaphragm muscle fibers converge and attach to the central tendon. During respiration, the cyclical cycles of muscle contraction cause the central tendon to shift from a relaxed to a taut sheet, causing changes in the volume and pressure within the thoracic cavity and promoting exchange of gases critical for respiration. To determine whether replacement of the central tendon with muscle might alter the elasticity and stiffen the diaphragm, leading to respiratory deficits, we assessed respiratory function by whole body plethysmography, which measures basal respiration in resting mice. We found that basal breathing rates and breath volume were not affected by these structural changes in the diaphragm (data not shown).

We reasoned that respiratory deficits may be unapparent during rest and only revealed during exercise. To assess respiratory function during exercise, mice were examined on a running wheel task, and the distance traveled over a 20h period was measured. We found that *abl2* mutant mice ran less than wild type littermates (Figure 5A). Because limb muscle development and limb strength were normal in *abl2* mutant mice (Figures S2 and Figure 5B), the decrease in running distance suggests that replacement of the central tendon with muscle in the diaphragm impaired respiration when challenged in an endurance task.

**Figure 5.**
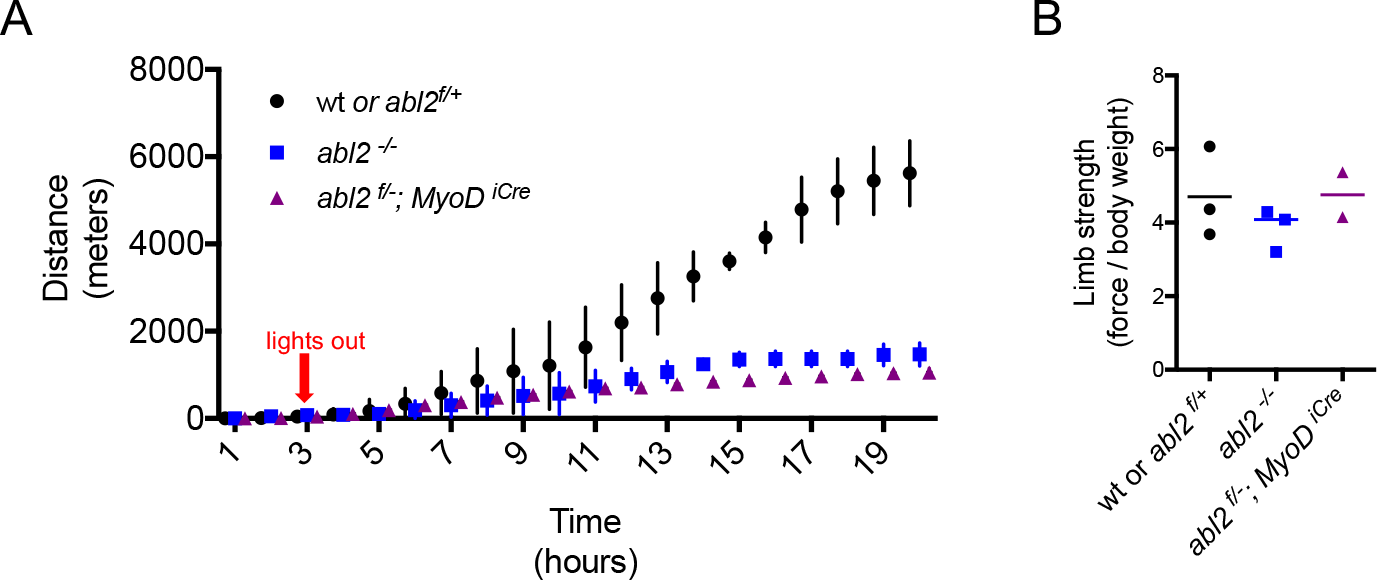
The exercise endurance of *abl2* null and *abl2* muscle conditional mutant mice is impaired. (A) During a twenty-hour period, *abl2*^-/-^ and *abl2*^*f*/-^ *MyoD*^*iCre*^ mice run less than wild type mice. (B) The limb strength of *abl2* mutant and wild type mice are similar. The graph and scatter plot show the values for individual mice and the mean values together with the SEMs. All *abl2* null and conditional *abl2* mutant mice were tested at 18 weeks with littermate controls.

*abl2* mutant mice display several behavioral abnormalities, including deficits in coordination, startle response and aggression, which are attributed to a role for Abl2 in the central nervous system (Koleske et al 1998). We therefore considered that a loss of Abl2 from the nervous system may be responsible for the reduced running performance of *abl2* mutant mice. To determine whether the reduced stamina of *abl2* mutant mice was due to a loss of muscle-derived Abl2 we studied the running performance of mice lacking *abl2* selectively in muscle (*abl2*^*f*/-^; *MyoD*^*icre*^ mice). Like *abl2* mutant mice, the *abl2* muscle-conditional mutant mice displayed reduced stamina on the running wheel (Figure 5A). Together, these results suggest that replacement of tendon with muscle in the diaphragm impairs respiration leading to impaired physical endurance.

We considered the possibility that the decreased endurance was due to a failure to develop slow, fatigue-resistant muscle fibers (Agbulut et al 2003). We therefore analyzed additional features of muscle differentiation in *abl2*^*f*/-^; *MyoD^icre^* adult diaphragm muscle and found that the distribution of fast and slow myofibers was normal (Figure S4). Therefore, the decreased stamina of muscle conditional *abl2* mutant mice cannot be attributed to a failure to develop slow muscle fibers. Further, muscle fibers in *abl2*^-/-^ adult mice displayed ultrastructural features characteristic of skeletal muscle, as the organization of sarcomeres and muscle-tendon attachment sites appeared normal (Figure S5). Thus, the decreased physical performance of *abl2* mutant mice is not caused by a failure to build the contractile apparatus and is more likely caused by the replacement of the elastic central tendon with less-tensile skeletal muscle.

### Abl2 negatively regulates myoblast proliferation

We hypothesized that the increased myoblast fusion in *abi2*^-/-^ mice might arise from an expanded pool of myoblasts produced by increased myoblast proliferation. To test this idea, we quantified the proliferation rate of myoblasts by measuring EdU (5-ethynyl-2’-deoxyuridine) incorporation in MyoD^+^ myoblasts in wild type and *abl2* mutant embryos. At E14, the percentage of myoblasts that incorporated EdU was greater in *abl2* mutant mice than wild type mice (Figure 6A,B), indicating that a loss of Abl2 leads to increased proliferation of myoblasts. Further, we measured the proliferation rate of myoblasts in primary cultures of diaphragm muscles from E18.5 *abl2* mutant and wild type mice. After growing cells at low density for 36h, the cultures were pulse-labeled with EdU for 1 hour, fixed, and stained for MyoD to identify proliferating myoblasts. We found that a higher percentage of *abl2* mutant myoblasts incorporated EdU than wild type control myoblasts (69.7 ± 4.1% and 44.1 ± 2.6% mean ± SEM respectively) (Figure 6C,D). In contrast, cells that did not express MyoD, such as fibroblasts or tendon cells, did not differ in EdU incorporation (Figure 6C,E). Thus, Abl2 negatively regulates proliferation of myoblasts but not of other cell types in muscle tissue. We also examined proliferation of myoblasts by double-staining with antibodies to MyoD and phospho-histone H3, a marker specific for cells undergoing mitosis, which revealed an increased number of mitotic myoblasts in *abl2*^-/-^ cultures compared to wild type littermate cultures (Figure 6F,G).

**Figure 6.**
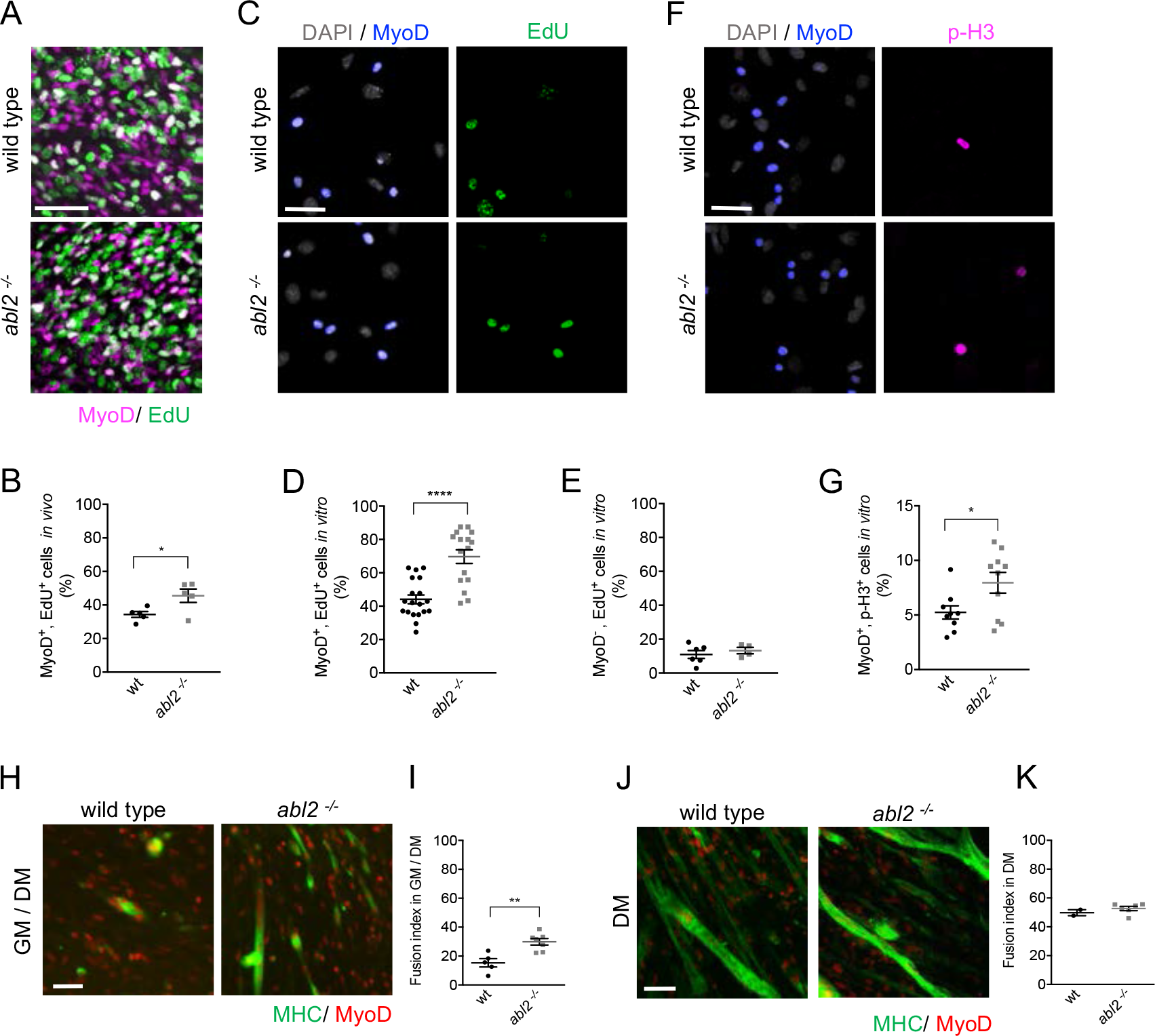
Myoblast proliferation is enhanced by a loss of *abl2*. (A,B) In E13.5-E14.5 diaphragm muscles in *vivo*, EdU incorporation is greater in *abl2*-/- myoblasts than wild-type myoblasts. (C,D) Cultured MyoD^+^ myoblasts from *abl2*-/- diaphragm muscles showed increased EdU incorporation. (C,E) In contrast, non-muscle cells (MyoD^-^) from *abl2* mutant and wild type mice proliferate at similar rates. (F,G) Staining for phospho-Histone H3 (pHH3), a marker for mitotic cells, was greater in *abl2* mutant than wild type myoblasts. (H,I) *Abl2* mutant myoblasts, which proliferated for 2 days before a switch to differentiation medium, displayed enhanced myotube formation. (J,K) *abl2* mutant myoblasts, which were plated at confluent density and directly into differentiation media, formed myotubes like wild type myoblasts. Scale bars are 50 μm in A,C,F,H and J.

To determine whether muscle fusion is enhanced by a loss of Abl2, primary muscle cells from *abl2*^-/-^ mice were propagated under proliferative conditions for 48 hours and then switched to differentiation medium for 48 hours. Myotube formation was greater in cultures from *abl2*^-/-^ than wild type mice, leading to a greater fusion index calculated by the ratio of fused myoblasts to total myoblasts (Figure 6H,I).

Although the increased proliferation of *abl2* mutant myoblasts could be solely responsible for the enhanced myoblast fusion, it remained possible that Abl2 also has a direct role in the mechanics of myoblast fusion. We therefore sought to determine whether the increase in myotube size was due solely to an increase in myoblast proliferation, thereby increasing the number of myoblasts available for fusion, or whether *abl2* mutant myoblasts also had an enhanced propensity for cell fusion. We plated primary cells at near confluent density into non-proliferative, differentiation conditions and scored for myoblast fusion two days later. Under these conditions, which are non-permissive for proliferation, wild type and *abl2*^-/-^ cultures had similar fusion indices (Figure 6J,K). Together, these data indicate that the longer myofibers in *abl2 ^/-^* mice arise from enhanced proliferation of *abl2^-/-^* myoblasts, increasing the size of the available myoblast pool, rather than an enhanced propensity of myoblasts to fuse.

Pax7-positive satellite cells are set-aside as a quiescent population of muscle stem cells during fetal development and are responsible for postnatal myofiber growth, homeostasis, and regeneration (Relaix et al 2005, Kassar-Duchossoy et al 2005, Keefe 2015, Lepper and Fan 2010). We wondered whether the expansion and fusion of myoblasts in *abl2* mutant mice might occur at the expense of establishing a Pax7-positive satellite cell population. We counted Pax7-positive muscle satellite cells in serial cross-sections of *abl2* mutant and control mice and found that Pax7-positive satellite cells were evenly distributed along the length of the diaphragm muscle fibers, and that total satellite cell number increased in *abl2*^-/-^ mice in proportion to the increase in muscle length (Figure S6). These findings indicate that increased muscle length did not occur at the expense of establishing the satellite cell pool and that Pax7-positive satellite cell numbers are established in proportion to muscle size.

### Ectopic muscle islands form within the central tendon of *abl2*^+/-^ mice

Mice that are heterozygous for *abl2* also display a muscle phenotype. In wild type mice, the central tendon of the diaphragm is devoid of muscle, whereas we observed ectopic muscle islands in the central tendon of *abl2*^+/-^ mice (Figure 7A). The number of ectopic muscle islands as well as the number of muscle fibers within each island were variable (Figure 7B), but individual muscle fibers within an island were uniform in length and aligned with respect to one another (Figure 7A). These findings suggest that the muscle fibers within an island coordinate their orientation and length, in a manner that is independent of contact with a preexisting specialized tendon cell border (Volk and VijayRaghavan 1994).

**Figure 7.**
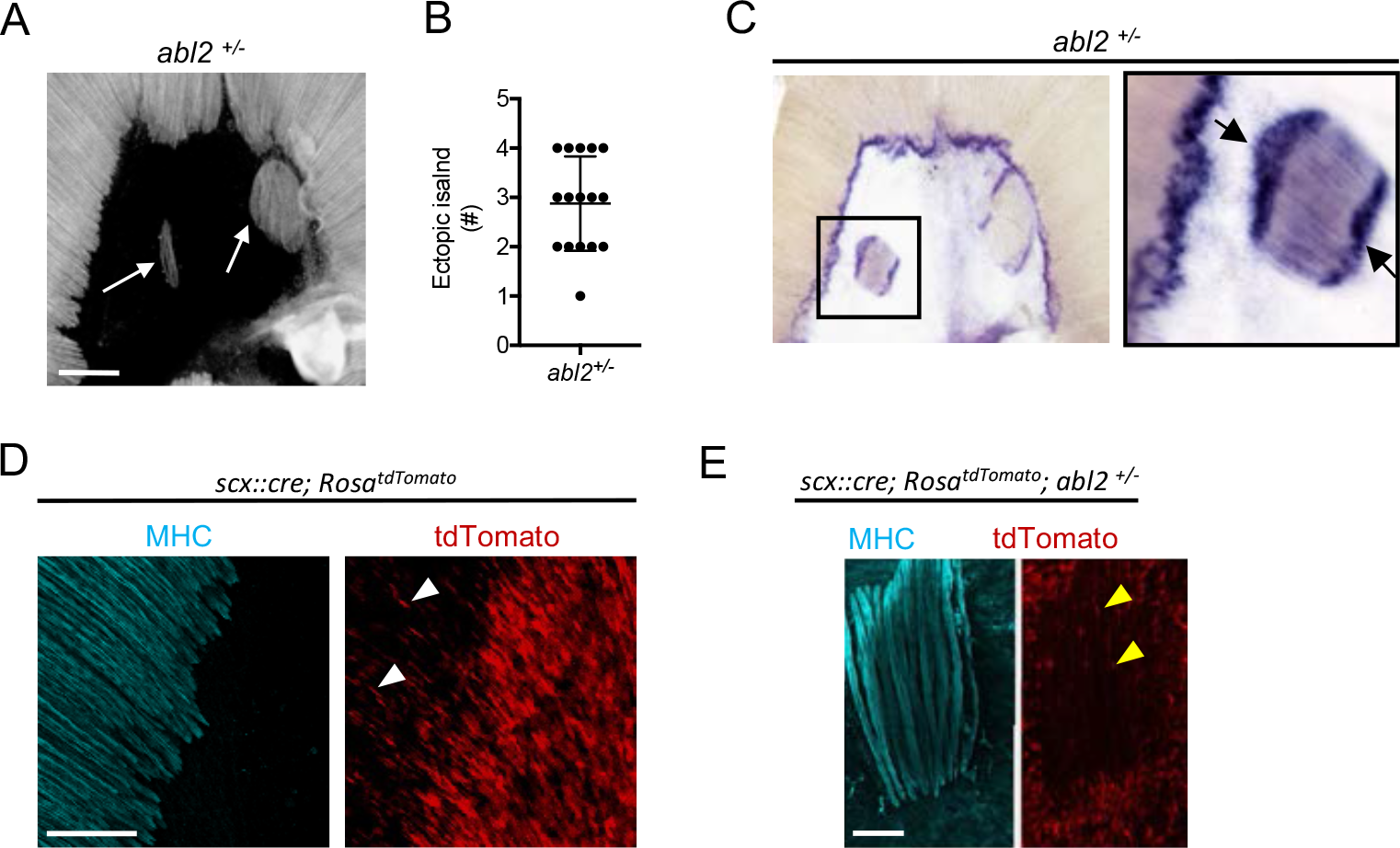
Ectopic muscle islands, which are found in the central tendon of mice that are heterozygous for *abl2*, induce tendon cell differentiation. (A,B) Whole mount diaphragms, stained with MHC, revealed ectopic islands in the central tendon of *abl2*^+/-^ mice. (C) *Scx* expression is enhanced at the ends of muscle fibers in the ectopic islands (black arrows) as well as at the normal MTJ in the diaphragm muscle. (D) Lineage-tracing experiments reveal tendon cells that intercalate between muscle fibers in the costal domain of muscle (white arrowheads). (E) In *abl2*^+/-^ mice, tendon cells intercalate between muscle fibers within the ectopic islands (yellow arrowheads) but do not contribute to myofibers. Scale bars are 500 μm in A, 250 μm in D, and 100 μm in E.

Although the location of the ectopic islands within the central tendon was variable, the orientation of the ectopic islands was biased to align with the nearby diaphragm muscle (Figure S7). This alignment of the ectopic muscle islands with the nearby diaphragm muscle raises the possibility that myoblasts, which are aligned with developing myotubes in the diaphragm muscle, retain this orientation as they aberrantly migrate, proliferate and differentiate in the central tendon. Alternatively, contraction of the diaphragm muscle may deform the central tendon in an oriented manner and thereby assist in aligning myotubes in the ectopic islands.

### Tendon cells become specialized when juxtaposed with ectopic muscle islands in *abl2*^-/-^ mice

We found that tendon cells at MTJs express high levels of *Scx* (Figure 2A). We examined whether tendon cells, positioned at the novel contact sites with the ectopic muscle islands, became similarly specialized. We found that *Scx* RNA expression was enhanced in tendon cells that are juxtaposed to the ends of the ectopic muscle islands (Figure 7C). These findings suggest that muscle fibers, even when displaced, provide signals to tendon cells, contributing to specialized features of tendon cells at MTJs.

### Tendon cell precursors do not contribute to muscle in ectopic islands of *abl2*^+/-^ mice

We wondered whether the muscle cells in the ectopic muscle islands formed from bona-fide muscle precursors or whether tendon cell precursors had altered their cell fate to form the ectopic muscle islands in *abl2*^+/-^ mice. We used *scx::Cre* mice and *Rosa*^*dTomato*^ reporter mice, which harbor a loxP-flanked STOP cassette preventing tdTomato transcription except in cells that express Cre recombinase, to trace the lineage of *scx*-expressing tendon precursor cells. We found that scx-expressing cells contributed to tendon cells surrounding and intercalated between muscle fibers but did not contribute to the muscle fibers within the ectopic muscle islands (Figure 7D). These findings are inconsistent with the idea tendon cell precursors switched their fate and converted to muscle to form the ectopic muscle islands in *abl2*^+/-^ mice.

## Discussion

We have uncovered a novel role for Abl2 in regulating muscle fiber length in mammals. We provide evidence that a loss of Abl2 function in myoblasts leads to enhanced myoblast proliferation and an expansion of the myoblast pool during embryonic development, leading to increased myoblast fusion and elongated muscle fibers.

It is striking that a loss of Abl2 leads to an increase in muscle length without a hypertrophic increase in muscle fiber diameter. Our *in vivo* and *in vitro* data describe an enhancement of myoblast proliferation and fusion during primary/embryonic myogenesis at E14 and during secondary/fetal myogenesis at E18.5 (Biressi et al 2007), which are developmental stages where myoblasts preferentially fuse to the ends of developing muscle fibers (Zhang and McLennan 1995). This mode of iterative nuclear accrual at ends of developing muscle fibers may favor longitudinal growth of muscle during embryonic development. It is possible that increased myoblast proliferation during early embryonic development favors longitudinal over hypertrophic muscle growth, whereas myoblast proliferation and fusion in adult myofibers may occur all along the myofiber and favor an increase in muscle fiber diameter.

Abl kinases are up-regulated in chronic myeloid leukemia, promoting cell proliferation. Abl2, however, attenuates cell proliferation in solid tumors, suggesting that Abl kinases can negatively or positively regulate proliferation, depending on cellular context (Greuber et al 2013, Gil-Henn et al 2013). Here, we provide evidence that Abl2 attenuates myoblast proliferation, limiting expansion of the myoblast pool and muscle growth. We do not know what lies upstream of Abl2 in myoblasts to control Abl2 activity, but it seems likely that multiple signals and steps modulate Abl2 activity to regulate myoblast proliferation and muscle growth.

Muscle precursors migrate to their final destinations using two distinct mechanisms. Body wall muscles, including the intercostal, vertebral, and abdominal muscles, extend and expand to envelop the body as a continuous outgrowth of the myotome (Christ et al 1983, Brand-Saberi and Christ 1999). In contrast, development of limb and diaphragm muscles requires delamination of muscle precursor cells from the dermomyotome, which engage in long-range migration to their target sites, where they proliferate, differentiate and fuse (Noden 1983, Dietrich et al 1999). Although intercostal and diaphragm muscles develop through different mechanisms, Abl2 has a role in governing muscle length in both muscles, indicating that Abl2 does not act primarily, if at all to regulate myoblast migration.

Abl2, however, does not regulate muscle fiber length in limb muscles. There is precedence for differing requirements for the development of diaphragm and limb muscles: although both limb and diaphragm muscles are derived from Lbx1-expressing precursors, migration of muscle precursors to the limb, but not the diaphragm, is perturbed in *Lbx1* mutant mice (Gross et al 2000). Thus, development of diaphragm and limb muscles differ in their requirement for Abl2, as well as Lbx1.

The role of Abl2 in some but not all muscles may be associated with the location, cellular environment or function of the muscle. Elongated muscle fibers are found in intercostal muscles, which attach to tendons that anchor the muscle to bone. Elongated muscle fibers are also found in muscles that are joined by tendons that interconnect two muscle segments, such as the central tendon of the diaphragm and the midline tendon between right and left levator auris longus muscles. However, muscles that attach to bone by force-transmitting tendons, such as those in the limb, remain unaffected by mutations in *abl2* (Murchison et al 2007).

In *Drosophila*, there is good evidence that muscle cells provide signals to tendon cells to induce their differentiation. Muscle cells produce Vein, a Neuregulin-like protein, which binds to the EGFR in adjacent tendon cells to mutually promote tendon cell differentiation (Yarnitzky et al 1997, Volk 1999). Activation of the EGFR in tendon cells enhances expression of Stripe and Slit, which signals back to Robo in muscle cells, contributing to muscle-specific tendon attachment (Kramer et al 2001, Volohonsky et al 2007). Although there is evidence supporting the idea that muscle and tendon cells signal to one another in vertebrates, these studies have largely focused upon TGF- and FGF-dependent signaling between the developing myotome and the adjacent sclerotome, where muscle is required for tendon formation (Brent et al 2003, Schweitzer et al 2010). Apart from this early axial signaling system, signaling between muscle and tendon in mammals has been not been fully explored. Unlike the formation of axial tendons, the induction of head and limb tendons does not depend upon muscle, although muscle and muscle contraction play a role in full tendon differentiation (Gaut and Duprez 2016).

We provide evidence that vertebrate muscle cells provide signals that are instructive for tendon cell differentiation, as myotubes that form as islands surrounded by tendon cells, induce tendon cell differentiation at the ends of the muscle fibers. Moreover, the myotubes within these islands are uniform in length and orientation, an unexpected finding if pre-positioned and pre-specialized tendon cells were required to organize muscle length and orientation. Instead, our findings raise the possibility that muscle pioneers, akin to those described in invertebrates (Ho and Goodman, 1983) may induce features of tendon cell differentiation, which subsequently lead to reciprocal signaling from tendon cells to muscle and contribute to an arrest of myotube growth, limiting myotube length and organizing the parallel alignment of myotubes.

We do not know whether enhanced myoblast proliferation plays a role in formation of the ectopic muscle islands in *abl2*^+/-^ mice. Muscle islands, however, are apparent in muscle conditional *abl2*^+/-^ mice (data not shown), suggesting that a decrease in Abl2 expression in myoblasts is responsible for formation of the ectopic islands. We considered the possibility that Abl2 might limit myoblast migratory behavior, preventing promiscuous migration of *abl2*^+/-^ myoblasts into the central tendon, but our cell culture studies failed to detect aberrant migratory behavior of *abl2* mutant myoblasts (data not shown). Nonetheless, the alignment of the ectopic muscle islands with the nearby diaphragm muscle raises the possibility that *abl2* mutant myoblasts, which migrated along the developing myotubes of the diaphragm muscle in a directed manner, aberrantly continued their oriented migration into the central tendon to form the ectopic muscle islands. In addition, myoblasts that inadvertently migrate into the central tendon domain may normally be eliminated, and this elimination process may be attenuated if Abl2 expression in myoblasts is reduced. Regardless of the mechanisms responsible for forming the ectopic islands, the organization of the myotubes within each island reveal mechanisms not previously understood and appreciated for aligning and setting the length of myofibers. Notably, although the muscle islands are enveloped by tendon cells, the myotubes are uniform in length and orientation, demonstrating that pre-positioned, specialized tendon cells are not essential to organize these features of muscle development. Myotubes may signal to one another to establish a common orientation and length. However, our results raise the possibility that specialized tendon cells, once induced by a developing, pioneer myotube, signal back to later arriving myotubes and contribute to their orderly arrangement.

## Materials and Methods

### Whole mount Immunohistochemistry

Diaphragm, internal intercostal and levator auris longus muscles were dissected from E18.5 embryos in oxygenated L-15 media, pinned onto Sylgard-coated dissection dishes, fixed for 1.5 hours in 1% para-formaldehyde (PFA) and blocked for 1 hour in 3% BSA (Sigma IgG free)+0.5% TritonX-100+PBS. Primary antibodies, diluted in 3% BSA+0.5% TritonX-100+PBS, were force-pipetted into the tissue. The muscle was incubated overnight on an orbital shaker in a humidified chamber at 4°C. Diaphragm muscles were washed 10 times over the course of 5 hours with 0.5% TritonX-100+PBS at room temperature and incubated in secondary antibodies, diluted in 0.5% TritonX-100+PBS, overnight on an orbital shaker at 4°C in a humidified chamber. Muscles were washed 10 times over the course of 5 hours with 0.5% TritonX-100+PBS at room temperature and rinsed in PBS before the muscle was whole-mounted in 50% glycerol. Muscles from at least 3 mice of each genotype were analyzed for each experiment.

### Cryosection Immunohistochemistry

Limb and diaphragm muscles were embedded in OCT media and frozen on a dry-ice platform. 10 μm sections, collected onto poly-L-lysine coated glass slides, were fixed in 1-4% PFA for 10 minutes, washed in 3% BSA+PBS 3 times for 5 minutes, permeabilized with 0.2-0.5% TritonX-100+PBS for 10 minutes, washed in 3% BSA+PBS and incubated overnight at 4°C with primary antibodies in 3% BSA/PBS (3% BSA+0.5% TritonX-100+PBS for Pax7) in a humidified chamber. Sections were washed in 3% BSA+PBS 3 times for 5 minutes before overnight incubation at 4°C with secondary antibodies, diluted in PBS, in a humidified chamber. Sections were washed 3 times for 5 minutes in 3% BSA+PBS, then PBS or counterstained with DAPI, before mounting in VECTASHIELD anti-fade mounting medium.

### Cell Culture Immunohistochemistry

Monolayer cell cultures were plated on Collagen-coated dishes, fixed in 1-4% PFA for 10 minutes, permeabilized with 0.2-0.5% TritonX-100 for 10 minutes and incubated with primary antibodies in 3% BSA+PBS overnight at 4°C in a humidified chamber. Cells were washed in 3% BSA+PBS 3 times for 5 minutes before overnight incubation in a humidified chamber at 4°C with secondary antibodies, diluted in PBS. Cells were washed 3 times for 5 minutes in 3% BSA+PBS, then PBS or counterstained with Hoechst 33342 before mounting in VECTASHIELD anti-fade mounting media.

### Single Fiber Dissociation

Diaphragm muscles were pinned onto Sylgard-coated dishes and digested with 2mg/ml Collagenase Type Ia (Sigma) in DMEM at 37°C for 30 minutes on an orbital shaker. After digestion, diaphragm muscles were flushed with 5% Fetal Bovine Serum in DMEM followed by DMEM buffered with 15mM HEPES. Diaphragm muscles were carefully dissected away from the central tendon and ribcage, and the myofibers were dissociated with fire-polished glass Pasteur pipettes under a dissecting microscope until individual fibers were released. Single fibers adhered onto poly-L-Lysine coated coverslips were fixed in 1% PFA for 5 minutes and washed in 3%BSA+PBS before overnight incubation in a humidified chamber at 4°C with primary antibodies, diluted in 3%PBS+PBS. Single fibers were washed in 3%BSA+PBS and incubated in a humidified chamber overnight at 4°C in primary antibodies, diluted in PBS. Single fibers were washed and counterstained with Alexa Fluor 594 Phalloidin (ThermoFisher Scientific) before mounting in VECTASHIELD mounting medium.

### C2C12 cell cultures

C2C12 cells were propagated in growth medium (GM: 10% Fetal bovine serum in Dulbecco’s Modified Eagle’s Medium supplemented with 4.5g/L glucose, L-glutamine and sodium pyruvate) at 37°C. Differentiation was induced, when cells were 80% confluent, by switching to DMEM supplemented with 2% heat-inactivated horse serum, 4.5g/L glucose and 1mM L-glutamine.

### Western Blotting and Immunoprecipitation

Cells were lysed in 30mM triethanolamine, 1% NP-40, 50mM NaF, 2mM NaV_2_O_5_, 1mM, Na-tetrathionate, 5mM EDTA, 5mM EGTA, 100mM N-ethylmaleimide, 50mM NaCl, 10uM Pepstatin A, Roche Protease inhibitor tablet (pH 7.4) for 30 minutes at 4°C. Lysates were centrifuged in a microcentrifuge for 20 minutes at 12,000 rpm at 4°C to remove cellular debris. Protein was measured by a Bradford assay, and samples were flash-frozen in SDS sample buffer or processed for immunoprecipitation.

Abl2 was immunoprecipitated from lysates with goat anti-Abl2 antibodies (Santa Cruz C20) and Protein-G Agarose Beads (Roche), following the manufacturer’s instructions. Protein was eluted from the beads with SDS sample buffer and fractionated by SDS-PAGE. Proteins were transferred onto PVDF membranes, which were probed with antibodies to Abl2 (Rabbit polyclonal, Proteintech). We used antibodies to pan-Cadherin (Cell Signaling Technologies) and Myosin heavy chain My32 (Sigma) to ensure equal loading of proteins in whole cell lysates.

### Primary cultures and *in vivo* Proliferation assays

Costal domains of diaphragm muscles were dissected from E18.5 embryos and separated from the central tendon in oxygenated L-15 medium under sterile conditions. Diaphragm muscles were minced and digested in 5mg/ml Papain, reconstituted in HBSS (+CaCl_2_ and MgCl_2_), for 30 minutes at 37°C with mild shaking. Digested cells were serially triturated with small bore fire-polished pipettes in 20% FBS in DMEM/F12 + 15mM HEPES media, passed through 40- μm cell strainers and plated at low densities (5.0 × 10^3^ cells per cm^2^) onto Collagen-coated dishes. After 36 hours in culture, cells were treated with EdU for 1 hour, following the manufacturer instructions, and subsequently fixed in 4% PFA, permeabilized in 0.5% PBS-TritonX100, processed for primary and secondary antibody staining and processed for Click-iT EdU Plus Alexa 488 imaging, following the manufacturer’s instructions (ThermoFisher Scientific). The scatter plots show the summed values from at least 150 cells in 5 random fields of view from individual cultures, as well as the mean ± SEM. For *in vivo* EdU analysis, pregnant mice, at 13.5-14.5 days of gestation, were injected IP with 0.5mg of EdU. Embryos were harvested 1 hour after *in utero* labeling; embryos were fixed with PFA, permeabilized in 0.5% PBS-TritonX100, stained with primary and secondary antibodies and processed for Click-iT EdU Plus Alexa 488 imaging, following the manufacturer’s instructions (ThermoFisher Scientific). Confocal images were captured from 3 regions of E13.5-14.5 diaphragms, collected from at least 3 embryos of each genotype: the myotendinous region of the left hemidiaphragm, the right hemidiaphragm, and the dorsal diaphragm.

### Primary Cultures and Differentiation Assays

Costal domains of diaphragm muscles were dissected from E18.5 embryos and separated from the central tendon in oxygenated L-15 medium under sterile conditions. Diaphragm muscles were minced and digested in 5mg/ml Papain, reconstituted in HBSS (+CaCl_2_ and MgCl_2_), for 30 minutes at 37^o^ C with mild shaking. Digested cells were serially triturated with small-bore fire-polished pipettes in 20% FBS in DMEM/F12 + 15mM HEPES media and passed through 40- μm cell strainers. Cells were re-suspended in 5% HS in DMEM and plated onto Collagen-coated dishes at 1.0 × 10^5^ cells per cm^2^. After differentiation, cells were fixed in 1% PFA, permeabilized with 0.5% PBS-TritonX100, and processed for primary and secondary antibody staining. Fusion indices were calculated by dividing the number of MyoD^+^ nuclei that are contained within MHC myotubes by the total number of MyoD^+^ nuclei.

### Imaging and data quantification

Images were acquired with a Zeiss AxioZoom stereo microscope and Zeiss LSM 700 or 800 confocal microscopes. Adjustments to detector gain and laser intensity were made to avoid saturated pixels. Muscle length, ectopic muscle island orientation, cross-sectional area was quantified using FIJI/ImageJ software. The data in scatter plots are expressed as mean values ± SEM unless otherwise noted in figure legends. Statistical difference between groups were evaluated using unpaired t-tests with Welch’s correction. *, p<0.05; **, p<0.01; ***, p<0.005; ****, p<0.001.

### *abl2* Riboprobes

Full length cDNA for *abl2* was purchased from OpenBiosystems (Accession number BC065912). 5’-ACACAGGCCTCCAGTGGG-3’ and 5’-GCACATCAGCTGGAGTGTGTTTC-3’ primers were used to amplify a 1088bp fragment, and 5’-AGGACCCTGAGGAAGCAGGGG-3’ and 5’-TCTGTGCCAATGAGCTGCACATC-3’ primers were used to amplify a 1321 bp fragment. Amplicons were subcloned into pBluescript KS plus vectors, digested with Kpn1/Hind3 and Sac1/Hind3 restriction endonucleases, respectively. DIG RNA labeling kit (Roche) was used to prepare digoxigenin-labeled anti-sense and sense strand riboprobes, following the manufacturer’s instructions.

### Embryonic Diaphragm Whole Mount *in situ* Hybridization

Diaphragm muscles were fixed overnight at 4° C in 4% PFA, dehydrated in methanol, treated with 6% H_2_O_2_ to inactivate endogenous alkaline phosphatase activity and digested for 30 minutes with 20μg/ml Proteinase K. Tissues were hybridized with digoxigenin-labeled riboprobes directed against mRNA encoding 1088bp and 1321bp fragments of *abl2* and processed using standard RNA *in situ* hybridization protocols (Kim and Burden 2008).

### Electron Microscopy

Adult diaphragm muscles were treated with 1μg/ml Tetrodotoxin and fixed in 1% glutaraldehyde. Fixed muscles where treated with 1% osmium for 1 hour, stained en bloc with saturated aqueous uranyl acetate for 1 hour, dehydrated, and embedded in Epon (Friese et al 2007).

### Behavior experiments

Running endurance was performed utilizing wireless running wheels and analysis software obtained from Med Associates Inc with age-matched female littermates at 18 weeks old maintained in a C57BL/6 genetic background. Mice were maintained in 12-hour light dark cycles and individual mice were placed in their home cage with one running wheel per cage, three hours prior to lights out on experimental days. Basal breathing analysis was performed using the Buxco Finepointe whole body plethysmography system. Grip force measurements were performed using a digital force gauge.

### Generation of *abl2* flox mice

Knockout first alleles of *Abl2* were generated as part of the International Knockout Mouse Consortium by IMPC/KOMP/EUCOMM (Skarnes et al 2011). Targeted ES cells were obtained and injected into B6-albino blastocysts. Chimeric mice were backcrossed to C57BL/6J mice and selected for germline transmission yielding “knockout-first” *abl2-tm1a* mice that possess an interrupting *lacZ* element, flanked by FRT sites, between exons 5 and 6 that serves as both a reporter and null allele. “Knockout-first” *abl2-tm1a* mice were bred to mice that express Flp recombinase under the control of *Pgk1* promoter (FLPo-10 Jax stock number 011065) to remove FRT flanked *lacZ* cassettes, yielding an allele that possesses LoxP sites flanking exon 6, e.g. *abl2* “flox” *abl2-tm1c* mice.

### Antibodies used for IHC/IFC

We used the following primary antibodies: Sigma mouse anti-My-32 (1:1000), Sigma mouse anti-NOQ7.5.4D myosin heavy chain slow (1:1000), Sigma Rabbit anti-Laminin (1:1000), Santa Cruz mouse anti-Pax7 DSHB bioreactor supernatant (1:500), Santa Cruz Rabbit anti-MyoD C-20 (1:1000), Thermofisher Mouse anti-MyoD 5.8A.

We used the following secondary antibodies: ThermoFisher Scientific Donkey anti-mouse (H+L) Highly Cross-Adsorbed Secondary Antibody Alexa Fluor 594 or Alexa Fluor 488, Donkey anti-rabbit (H+L) Highly Cross-Adsorbed Secondary Antibody Alexa Fluor 594 or Alexa Fluor 488, Goat anti-Rat IgG (H+L) Cross-adsorbed Secondary Antibody Alexa Fluor 488, Life Technologies.

## Acknowledgments

We are grateful to Dr. Sang Yong Kim, director of the Rodent Genetic Engineering Core at NYUMC, for his assistance in deriving the *abl2*^*flox*^ mice described here. This work was supported with funds from the NIH (NS36193 and NS075124). We thank Maartje Huibers and Julien Oury for their comments on the manuscript.

---

**Figure S1.**
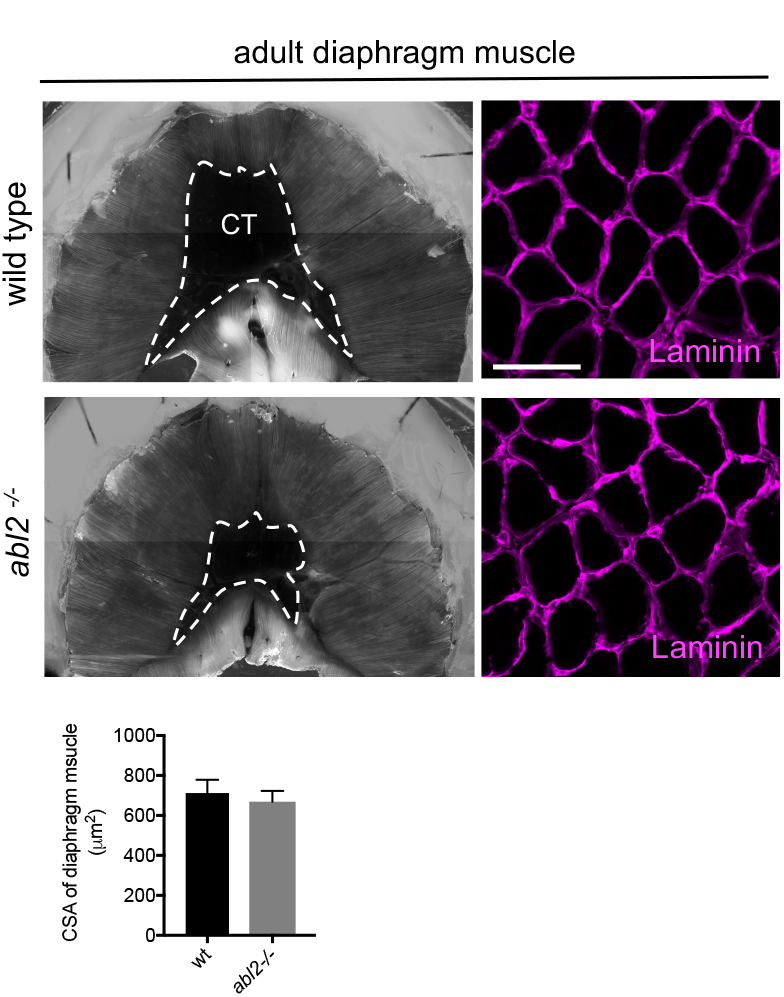
*abl2*^-/-^ mice retain muscle phenotypes throughout adulthood. Diaphragm muscle fibers are excessively long in adult *abl2*^-/-^ mice. The cross-sectional area of the *abl2*^-/-^ myofibers, however, remained normal, without signs of hypertrophy or atrophy. The central tendon (CT) is denoted by dashed lines. The means ± SEMs for three animals in each group are shown. Scale bar is 50 μm.

**Figure S2.**
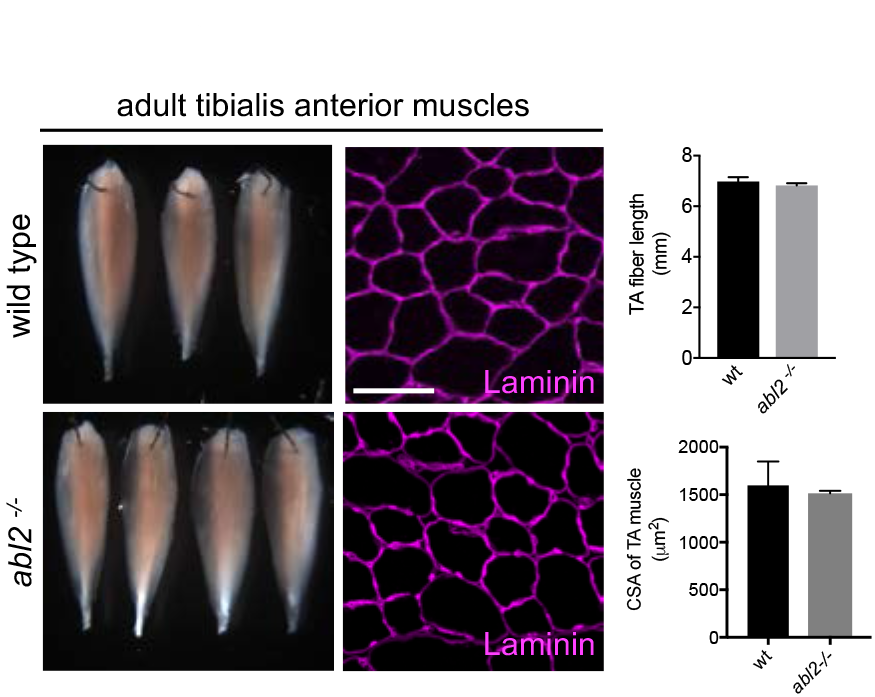
Limb muscles appear normal in muscle length and cross-sectional area. The length and crosssectional area of *abl2*^-/-^ tibialis anterior (TA) muscles are normal. The means ± SEMs from the left TA muscles of 3-4 mice in each group are shown. Scale bar is 50 μm.

**Figure S3.**
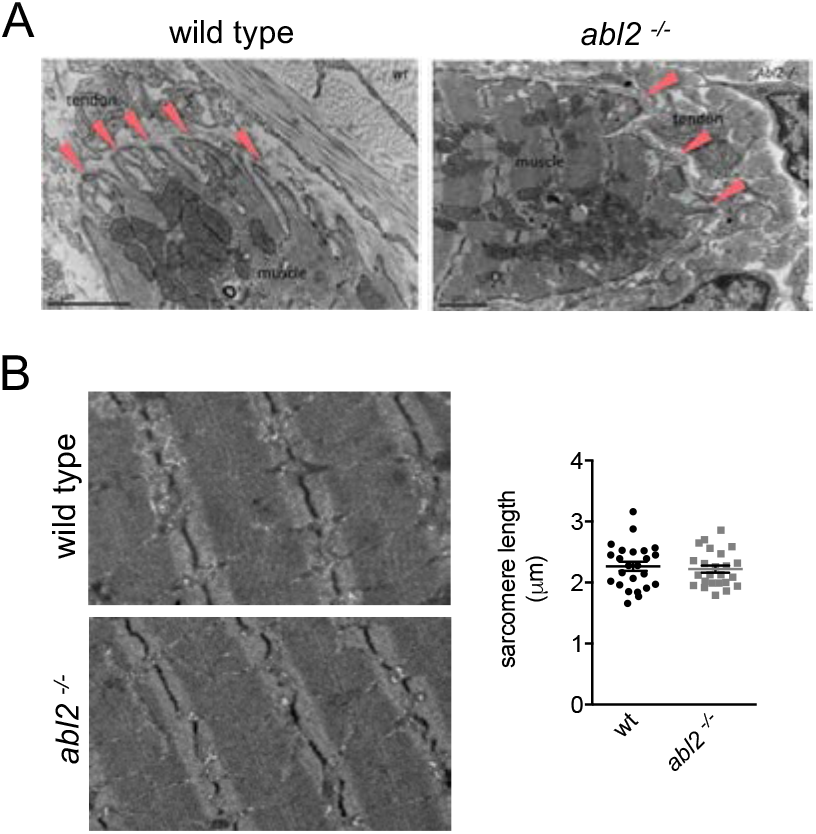
The ultrastructural of the myotendinous junction and sarcomere appear normal in *abl2*^-/-^ mice. (A) Deep muscle membrane specializations form at the myotendinous junction in wildtype and *abl2*^-/-^ mice. (B) The scatter plot shows the sarcomere lengths for 12 different fibers from each of two wildtype and *abl2*^-/-^ mice. The mean ± SEMs are also shown.

**Figure S4.**
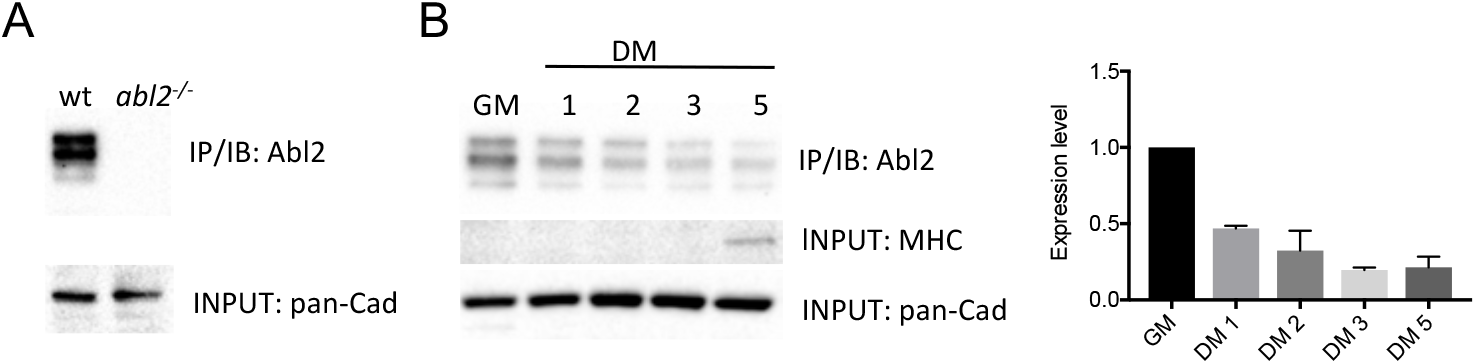
Abl2 expression is enriched in proliferating myoblasts. (A) Cadherin levels (pan-Cad) were measured in cell lysates, and Abl2 was immunoprecipitated (IP) from lysates containing an equal level of pan-Cad in wild type and *abl2*^-/-^ primary cultures to validate the IP and immunoblotting (IB) antibodies. (B) Abl2 was immunoprecipitated from lysates of C2C12 cultures grown in growth medium (GM) or in differentiation medium (DM) for the indicated number of days. Abl2 is highly expressed in C2C12 cells in growth media as 3 distinct bands between 100-150 kDa, and expression declines during differentiation. Abl2 expression in GM was assigned a value of 1.0, and all other values are expressed relative to GM.

**Figure S5.**
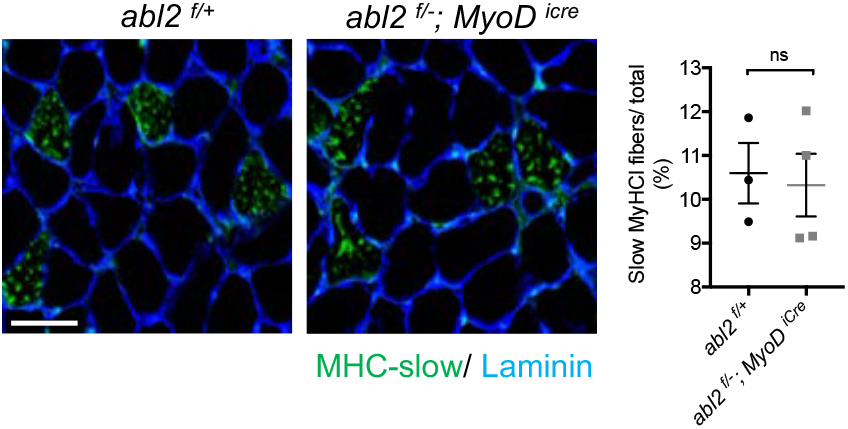
Adult diaphragm muscles do not exhibit a change in slow fiber-type number. 150-300 total myofibers were examined in crosssections from each mouse. The scatter plot shows the percentage of myofibers that are stained by antibodies to slow MHC. The means and SEMs from three control and four muscle-conditional *abl2* mutant mice are shown. Scale bar is 50 μm.

**Figure S6.**
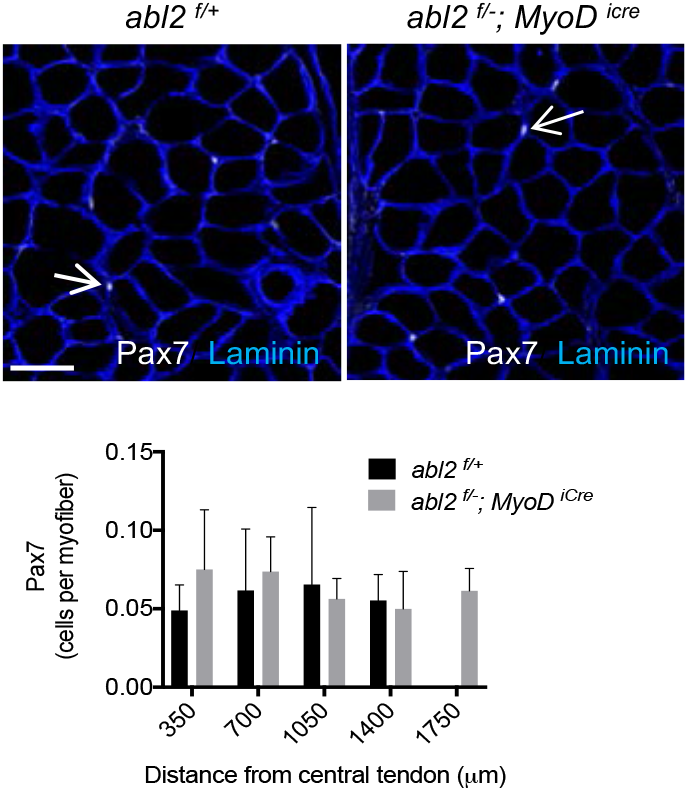
The number of Pax7^+^ cells is increased in proportion to the increased length of myofibers. Longer muscle fibers in *abl2*^-/-^ mice have proportionally more Pax7^+^ cells. The graph shows the mean (± s.d.) number of Pax7^+^ cells in serial cross sections from three mice in each group Scale bar is 50 μm.

**Figure S7.**
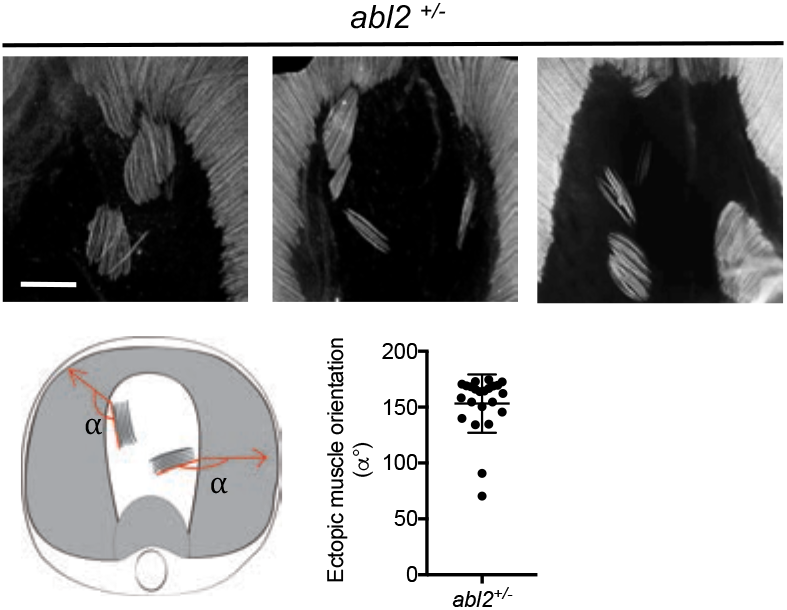
The orientation of ectopic muscle fibers correlates with the orientation of the main costal diaphragm myofibers. The scatter plot shows the angle of each ectopic island in relation to the orientation of the myofibers in the nearest costal diaphragm muscle. The mean value and ± s.d. are shown. Scale bar is 500 μm.

